# A Vector Navigation and Inference Architecture can Construct Universal Cognitive Maps for Abstract Reasoning

**DOI:** 10.64898/2026.03.07.705326

**Authors:** Andrej Bicanski

## Abstract

The idea of cognitive maps in the hippocampal formation has its origins in decades of spatial cognition research. However, increasing evidence points to cognitive maps also supporting non-spatial tasks. It appears that the hippocampal-entorhinal system can map any combination of systematically varying stimulus dimensions onto the same neural manifolds that support spatial representations - grid cells. Here I propose a model that shows how spatial navigation architectures can iteratively construct universal cognitive maps. The model is neurally plausible and exhibits noise-tolerance that implies specific behavioral predictions. The mapping process accommodates discontinuous stimulus spaces and the model can be used to support several types of abstract reasoning, including analogy making, subspace construction, and perspective taking. Importantly, these capabilities are supported by the same neural processes that are used to build a map. Thus, the combination of mechanisms at play suggests how spatial navigation architectures can function as domain-general substrates of cognition.

## Introduction

Since Tolman’s original proposal^1^, the concept of the cognitive map has first been fleshed out in the context of spatial memory and navigation^2^, principally due to the discovery of various spatial neural codes^3–8^. More recently experiments have expanded the notion to other domains of cognition^9–14^, suggesting that the hippocampal-entorhinal system can map arbitrary stimulus dimensions on low-dimensional neural manifolds in entorhinal cortex^15–17^. In humans, such maps are taken to be indicated by six-fold modulation of the BOLD signal, inherited from the neurons that instantiate the map - grid cells^4,18^. In spatial paradigms, grid cells exhibit firing locations on a regular hexagonal grid, at varying scales, which has led to various proposals about their role in spatial memory and (vector) navigation^19–26^, as well as imagination^27–29^.

Computing a shortcut, i.e., a vector across unexplored terrain, can be viewed as an inference in the spatial navigation domain. Analogously, a vector from point A to point B on an abstract two-dimensional plane, spanned by arbitrary stimulus dimensions, constitutes an inference about the relation between the conceptual entities anchored to A and B. How the anchoring of such stimuli can be accomplished in a neurally plausible way is unknown. Here it is shown how such universal cognitive maps (UCMs) can be produced by adapting spatial navigation architectures that depend on grid cells^4^. To repurpose nominally spatial grid cells is reminiscent of a long-standing notion: that the hippocampal-entorhinal declarative memory system can be unified with the neural infrastructure for spatial navigation^30,31^. Here this notion is further extended.

The present model is applied to two datasets. First, public data from the Open Affective Standardized Image Set (OASIS)^32^, a database of images with valence and arousal ratings, previously used to demonstrate non-spatial grid cells^33^ for an affective state-arousal cognitive map. Second, a realization of “bird-space”, which provided the first evidence for abstract grid maps^9^. The algorithm for constructing UCMs relies on three components. First, inputs are mapped by leveraging similarity along isolated stimulus dimensions, which is taken to be directly proportional to a distance metric^34^ provided by grid cells. Second, a vector navigation architecture (VNA). Third, a positional inference network (PIN). The VNA can retrieve vectors between points, while the PIN can infer the target position from a starting point and a vector. Beginning with an anchor (the first stimulus on the map), the PIN can iteratively populate the metric map. Following the construction of the UCM, the VNA and PIN can be combined to achieve reasoning capabilities, such as analogy formation, subspace construction, and egocentric perspective taking in the map.

## Model outline

Real-world stimuli vary along numerous dimensions. We assume that there is a record of the magnitudes of individual stimulus attributes upstream of hippocampal-entorhinal system (see Discussion for a proposal). Crucially, these magnitudes are initially not related to each other. The UCM algorithm operates as follows. The first stimulus is anchored to an arbitrary position on the grid map (potentially mediated by hippocampal indices^3,35^). The exact location on the map is irrelevant given the periodic structure of the toroidal grid cell manifold coupled with large capacity^19,23,26^. The second stimulus is placed at a distance (on the grid metric) proportional to the similarity to the first stimulus (along the relevant dimensions). The proportionality factor is equivalent to the velocity gain in biologically plausible models of grid cells^20,21^, which determines how far the pattern of neural activity on the toroidal grid manifold translates for a given real-world distance (time-integrated velocity input). Crucially, this gain is known to be plastic^36,37^, which we exploit as a key premise here. That is, the scale can be set and potentially adapted further at a later time. After more than two stimuli have been observed, a coherent cognitive map can be constructed iteratively via triangulation. Figure 1 outlines the complementarity of VNA and PIN, the mapping algorithm and two possible neural implementations (Figure S1 illustrates the mapping procedure for the first 4 stimuli). Locations on grid map correspond to population vectors (henceforth PVs) across the grid cell network (Figure 1a). A VNA can compute a vector between two PV-coded locations A and B. However, the complementary computation, directly inferring B given A and the displacement vector, has not been explored before (Figure 1b-d). Such a network can directly output the grid cell PV of the location at which the next stimulus should be anchored on the grid map (Figure 1e). Here, a distance-cell based architecture (cf. ref.^26^) is employed since it requires fewer computational steps, but other neural implementations, e.g. via sweeps through the head-direction network (cell signaling orientation in spatial settings), could also be employed (Figure 1f-h).

**Figure 1:**
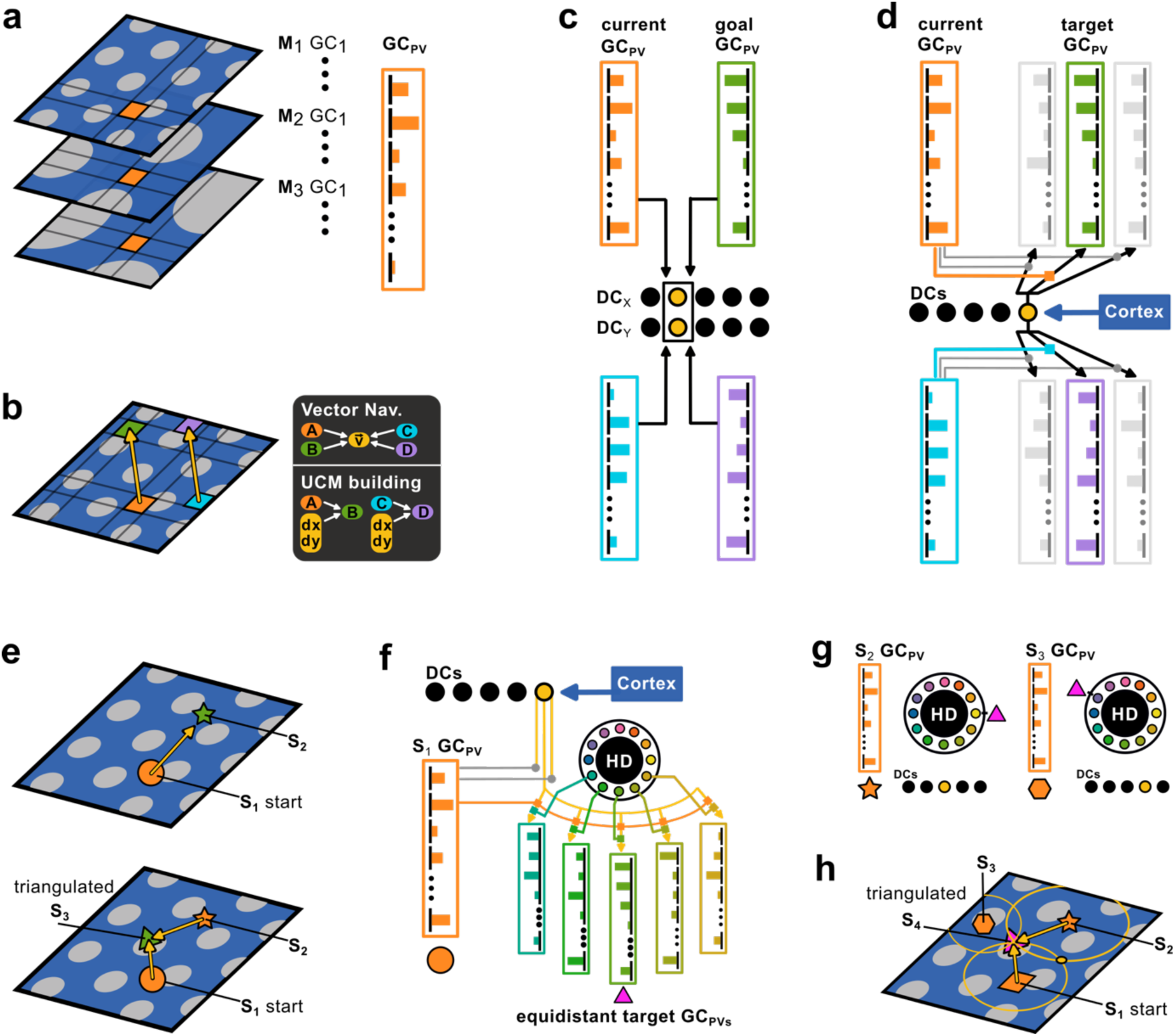
Construction of UCMs as a process complementary to vector navigation. **(a)** Illustration of stacked grid cell firing rate maps across modules (M_1_, M_2_…) and a grid cell (GC) population vector (PV) at a given location (orange square). **(b)** Vector navigation requires the computation of relative vectors (v) between locations A, B, or C, D, given by grid cell PVs. Building a UCM requires inferring location B given point A and a displacement vector. **(c)** Simplified illustration of a distance cell (DC) network in which multiple pairs of grid cell PVs activate the same distance cells along two dimensions (nominally x, y) to compute the yellow vectors in (b). **(d)** A positional inference network (PIN) can act complementary to vector navigation, target grid cell PVs can be inferred if the orange grid cell PV can modulate (e.g., via pre-synaptic gating) the activation (provided by distance cells) of the green grid cell PV. In this case, distance cell activation is assumed to be supplied by cortical computations (see main text). **(e)** UCM construction process: the PIN (d) can triangulate coherent map positions on the grid sheet as follows. Top panel: Stimulus S_1_ (orange circle) is anchored to an arbitrary position in grid space. Stimulus S_2_ (green star) is anchored at the correct distance from the orange circle, in an arbitrary direction. Bottom panel: subsequent stimuli (S_3,4,5_ …) can be placed at overlapping candidate locations that respect the cortical distance measures, computed from 2 or more anchors (already placed stimuli, here S_1,2_, etc.). Also see Figure S1. **(f)** An alternative algorithm for triangulation can be implemented if head direction cells as well as the current position grid cell PV (orange) modulate the activation of target grid cell PVs. See squares on connections from distance cells to all grid cell PVs that represent the set of equidistant (in all directions) positions relative to the current state. **(g)** Grid cell PVs for different (already placed) stimuli (here star and hexagon, see (h)) are at different distances (see DCs) and directions (see HD ring) relative to the magenta triangle – which is the stimulus that is to be placed on the grid map. **(h)** Two anchored stimuli would leave two possible placements for the magenta triangle (see triangle and small yellow circle). With three or more stimuli already anchored, the intersection of circles (HD sweeps) yields unambiguous triangulation.

### Constructing UCMs from Datasets

The first dataset consists of scores of emotional valence and arousal – the OASIS dataset^32,33^. These scores already represent a magnitude code for two individual dimensions. Though the data can be visualized in a 2D scatter plot, the mapping on a 2D neural representation remains to be accomplished. Exploration of how a magnitude code might form is relegated to the Discussion. Figure 2a shows representative examples of grid cell firing rate maps (as approximation to a fully dynamic, toroidal treatment) across 6 modules of grid cells at different scales. Each plot shows one representative cell from a module with 100 neurons each. The scale is arbitrary given grid spacing is plastic^36,37^, which is a key premise of the present model, and is compatible with biologically detailed grid cell models. This variable gain serves as the proportionality factor between grid scales and the magnitudes in stimulus space. To approximate this gain-setting, the valence and arousal magnitudes of the OASIS dataset are rescaled to approximately the range of the grid cell rate maps (Figure 2c,d). A subset of data points is employed, similar to experiments^33^. Figure 2b re-iterates the complementarity of the PIN and VNA in the visual domain^12,38^. Having rescaled the data, we can employ the algorithm outlined above to construct the valence-arousal map. Each point in the dataset corresponds to valence and arousal ratings for a specific stimulus and it is these stimuli that are to be anchored the grid map. To build the UCM, a first valence-arousal point (i.e., image, stimulus S1) is chosen and arbitrarily associated with the grid map (Figure 2d). Then a second stimulus S2 is chosen. The difference in arousal and valence rating between S1 and S2 is taken to represent the vector from S1 to S2. That is, (dis)similarity is quite literally distance^34,39,40^. Armed with the displacement vector and grid cell PV for S1, the PIN can calculate the grid cell PV at which to anchor stimulus S2. Successive stimuli are then anchored iteratively, by inferring their grid cells PVs from at least 2 anchors (also see Figure 3 below).

**Figure 2:**
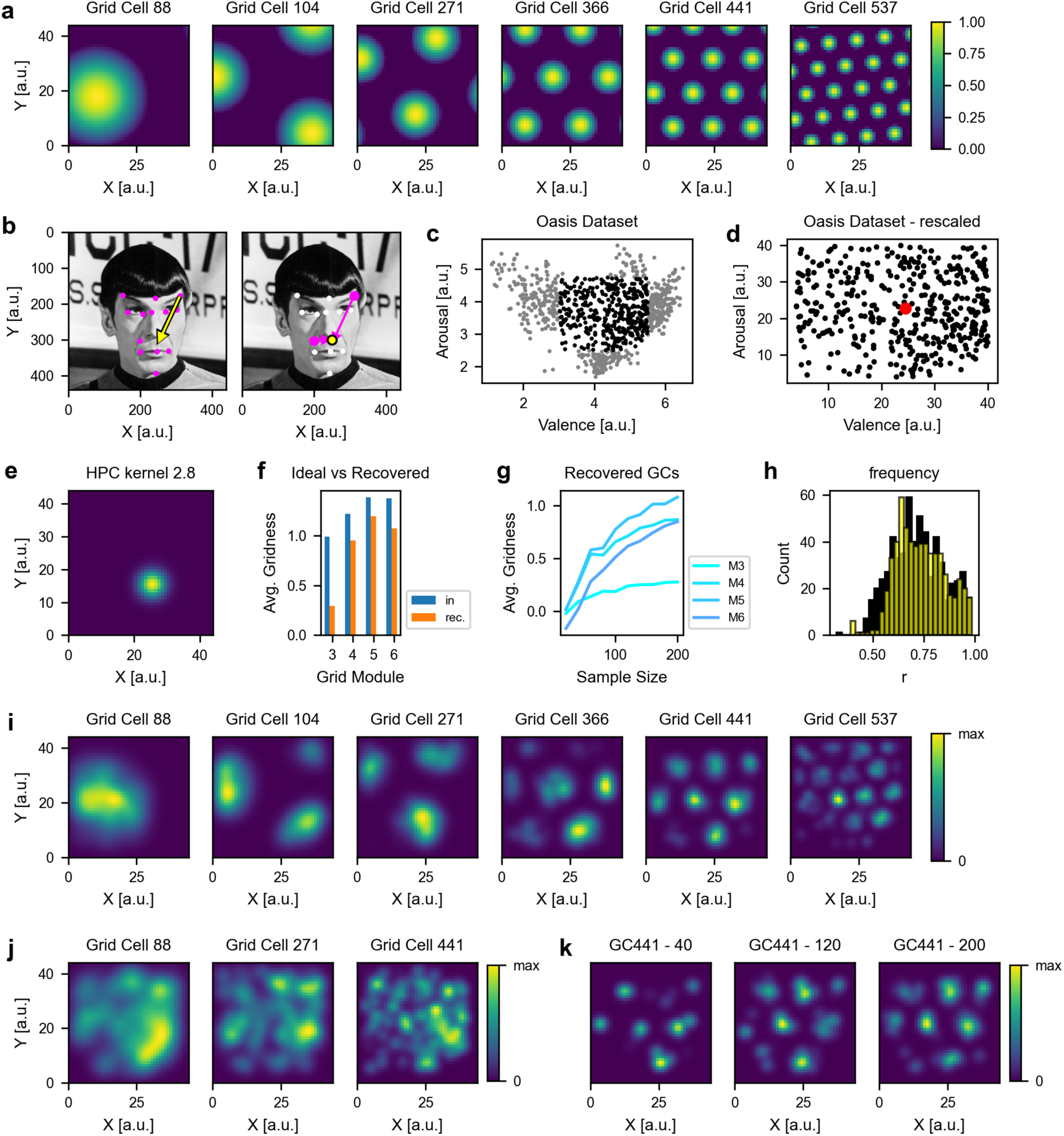
A cognitive map of valence and arousal constructed via location inference. **(a)** Example grid cells from six modules of input grid cells. **(b)** The visual (eye-movement navigation) analogy (cf. ref.^38^). Magenta is given, yellow is inferred. In a VNA the points are given (left) and the displacement vector is inferred (left). Here, the PIN is supplied with a starting location and displacement vector (magenta arrows, two inferences towards the same target) and infers the target grid cell population vector (PV) at the location marked by the yellow dot (right). c) The oasis dataset, and d) the reduced dataset rescaled to the dimensions of the grid cells, assuming plastic scaling. Red dot indicates first stimulus. e) A Gaussian kernel akin to a place cell used to multiply PVs to estimate grids from mapped stimuli and memorized PVs (see main text), here sigma 2.8. f) Gridness calculated from autocorrelograms of reconstructed grid cells vs from input grid cells. Modules M1 and M2 do not show hexagonal structure in autocorrelograms due to large scale. g) varying the number of samples used for reconstruction/mapping, it can be seen that gridness is high already with low sample numbers for modules with sufficient peaks for gridness analysis. h) low gridness does not automatically imply lack of grid coding, as evidenced by high correlations between recovered grid cells and input grid cells on a cell per cell basis (OASIS dataset in black, surrogate data with 10-fold more data points in yellow). High correlation bins include cells with low gridness due to scale. i) Recovered grid cells. j) Shuffling vector distances used in reconstruction breaks grid structure. k) Evolving data grids with different sample sizes (of valence-arousal stimuli) used to construct them (here 40, 120, 200). Already at low sample size the grid structure is evident.

**Figure 3:**
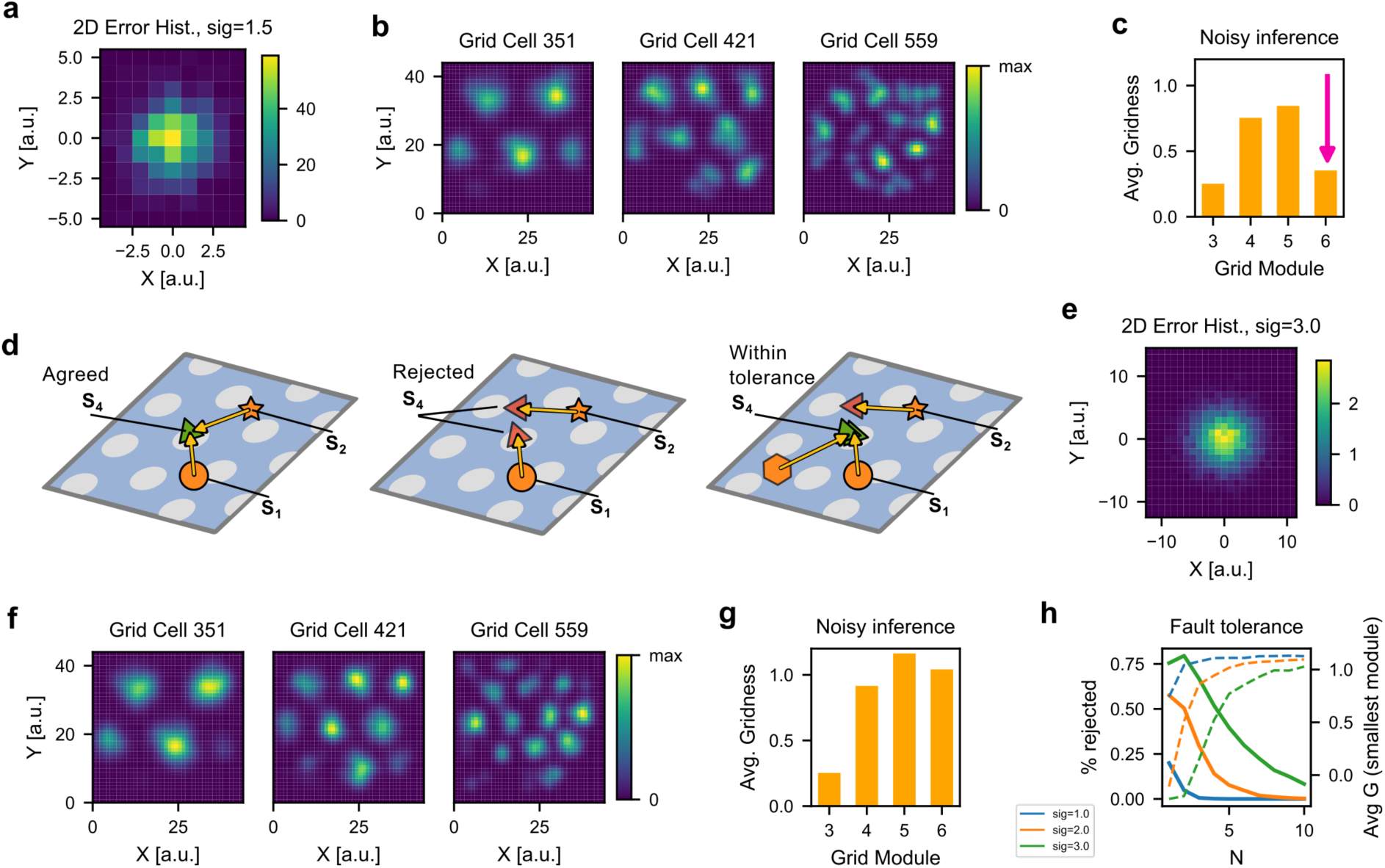
UCMs can be fault tolerant at the expense of task time. **(a)** 0-5% percent noise in valence-arousal vectors leads to **(b)** small grid cells becoming non-recoverable when each stimulus is triangulated only once. **(c)** Average gridness across modules under noise conditions. **(d)** A solution to noise in the system. When positional inference fails to yield agreement in grid cell PVs (coordinates on the grid map) from two anchors, we reject placement of the stimulus on the grid map (middle panel). Recruiting multiple anchors yields fault-tolerance (right panel, green estimates with tolerance, red estimate too far from the mean). **(e)** Large noise (up to approximately 10% grid map range, cf. panel a) can be accommodated by the correction mechanism. I.e., it is required that multiple attempts at inference yield nearby coordinates on the grid sheet (i.e., similar grid cell PVs). **(f)** Resultant dataset grid maps recover smaller scales. **(g)** Average gridness in the smallest module is restored, despite increased noise. **(h)** The number of unanchored stimuli (solid lines) and avg gridness (dashed lines) in the smallest grid scale as a function of the number of triangulations for three noise levels

To test if the valence-arousal space has been adequately mapped onto the grid cell network is to test that each valence-arousal-pair is associated with a unique grid cell PV, such that their relative distances are preserved on the grid metric. To read out this grid structure we can multiply a given stimulus-specific grid cell PV with a Gaussian kernel centered on the coordinates of a stimulus in re-scaled stimulus space (Figure 2e, Methods). To build intuition for the Gaussian kernel, let us assume that each grid cell PV is re-activated by a cell that also connects to the distributed stimulus representation upstream of hippocampus, akin to a hippocampal index. These same indices could also associate a stimulus with additional landmarks. As a stand-in for re-activation of the grid cell PV by a hippocampal index we multiply the memorized grid cells PV with the place cell-like Gaussian kernel. However, this kernel has to be placed at ground truth stimulus space coordinates to allow for potential mismatch between the output of the PIN and ground truth relations (see noise simulations below), and to mimic binning of PVs by an experimenter. Repeating the process for all stimuli and summing up their contributions should recover the grid structure if, and only if, mapping was successful. Figure 2i shows the recovered grid cells. These recovered grid cells can be considered a proxy for what one would record if one had access to single cell activity. They are not intermediate stages in some learning process, because we are repurposing existing spatial grid cells in the mapping process. Figure 2f shows the average gridness calculated from autocorrelograms (a measure of hexagonal symmetry for grid cells, see Methods) across modules, comparing input grid cells (Figure 2a) with recovered grid maps (Figure 2i). Large modules do not show hexagonal symmetry in autocorrelograms due to few firing peaks (see cell 88 and 104 in Figure 2a,i, also see supplementary Figures S2.1, S2.2), but correlating input grid cells with recovered grid maps on a cell-by-cell basis shows high overlap (Figure 2h). Surrogate data confirms this trend (Figure S2.2). Together with visual inspection this is taken to confirm that the OASIS stimuli have indeed been correctly mapped to appropriately spaced grid cell PVs. When vectors between points in arousal-valence space are scrambled, the recovered grid cells lose their hexagonal firing structure (Figure 2j). Finally, the grid structure in the recovered data grid cells is apparent with as little as 50 or fewer OASIS stimuli (Figure 2g) out of the subset of 409. That is, the structure of the UCMs emerges quickly under favorable conditions (cf. to noise simulations below). Figure 2k shows the evolution of a representative cell. Note that the fact that early data grids don’t look like fully formed grid cells at low sample numbers is simply due to not having accumulated enough activity to make a good rate map, further highlighting that these recovered grids should be considered a proxy for experimental recordings. If the inference process were flawed, the final pattern would diverge from a hexagonal grid.

The iterative construction of UCMs, based on triangulation from multiple anchors has a notable consequence. For Figure 2 we engage the PIN from two different anchor positions (already mapped stimuli) to infer location for each target. If two such inferences fail to point to similar grid cell PVs (within some tolerance), we can use this failure to signal errors. This endows the model with fault-tolerance and yields specific behavioral predictions. To test fault tolerance, Gaussian noise was added to the vectors between anchors and targets in valence-arousal space (Figure 3a). To a first approximation this noise can be taken to represent the combined effect of noisy arousal-valence differences, PIN imprecision, and noise in the grid cell representation. The effect on recovered data set grid maps is most pronounced in small modules (Figure 3b) and is reflected in a drop in gridness for that module (Figure 3c). However, larger modules still exhibit grid patterns, albeit with reduced gridness (Figure 3c,g), suggesting that noise (here on the order of 5% of the grid cell coordinate range, cf. Figure 3a) does not catastrophically impede grid mapping within some resolution. Rather, the system degrades progressively, losing the smallest scale first. Figure 3d outlines the solution to noise: we can increase the number of anchors in the algorithm. Each anchor yields inferred target coordinates/PVs and we can place the target stimulus where we find most overlap. That is, noise can be compensated simply by increasing the number of triangulation events (attempts at inference). Note that this does not imply more observations. Instead, more anchors must be retrieved from memory. This could happen during a task, or potentially later, via memory replay (see Discussion). Overlap could be defined by the mean, median or by taking a majority vote across inferences to settle on a target grid cell PV. Here the average of coordinates within some distance from the mean (tolerance) is used for readout (see Methods). If all inferred coordinates fall outside the tolerance we refuse to anchor the stimulus to the grid map. This procedure can deal with high noise (Figure 3e, up to 10% of grid range), and recovers the smallest grid scale (Figure 3f,g, Figure S3), albeit at the expense of more attempts to infer locations. The number of unanchored stimuli drops quickly with more anchors while gridness of the smallest module increases (Figure 3h). That is, the model algorithm is consistent with the notion that in experiments “high noise” (i.e., a difficult task) requires more trials to learn the 2D embedding. Too difficult a task may never yield a grid map in the time allocated to the task as overlap/majority votes become rare and no stimuli are anchored. Though not used here, an additional noise-compensation mechanism would consist in checking the output of the PIN by inserting the inferred goal location and the start location PVs into the VNA to check if the vector it yields is similar to the one read from the (rescaled) dataset. A discrepancy could trigger a new attempt at mapping. Similarly, we could map each stimulus multiple times and take its average inferred position. This highlights that the interplay of VNA and PIN opens up several possibilities for dealing with noise.

The second dataset used here is a synthetic variant of the “stretchy birds” stimuli, used in the seminal study of Constantinescu and co-workers^9^. Birds were procedurally generated to cover a range of leg and neck lengths (Figure 4a-c), the two dimensions which are to be mapped. A simple linear decoder architecture was used to get neck and leg length back. While this is not strictly necessary to get the lengths out of the dataset, it acknowledges that some upstream brain areas (relative to entorhinal cortex grid cells) must be capable of extracting the magnitude of individual stimulus features (see Discussion below). Like for the OASIS data set, it was assumed that the plastic gain on grid scales can ensure that the observed numerical leg and neck length range can be scaled to adequately leverage the range of the grid map. Following rescaling (Figure 4a) moderate noise was added on top of the dataset (Figure 4b). Data points cluster around their generated classes, different from the OASIS dataset. Then – employing the algorithm described above – bird space is mapped onto the grid cell network via the PIN, with each bird anchored to a specific grid cell PV. The recovered grid activations (Figure 4d,e) show prominent hexagonal patterns, high average gridness (Figure 4f) and strong correlations with input grid cells (Figure 4g), confirming that the positional inference network successfully anchored all bird stimuli to the grid maps. Patterns are slightly more pronounced compared to the OASIS dataset because more stimuli are used. The fact that birds cluster and are not uniformly distributed does not impede the mapping. Finally, bird space allows for an illustrative visualization. Figure 4h displays individual birds linked to the location at which they were encoded on an individual, representative grid cell. It can be seen that leg length and neck length vary consistent with map position.

**Figure 4:**
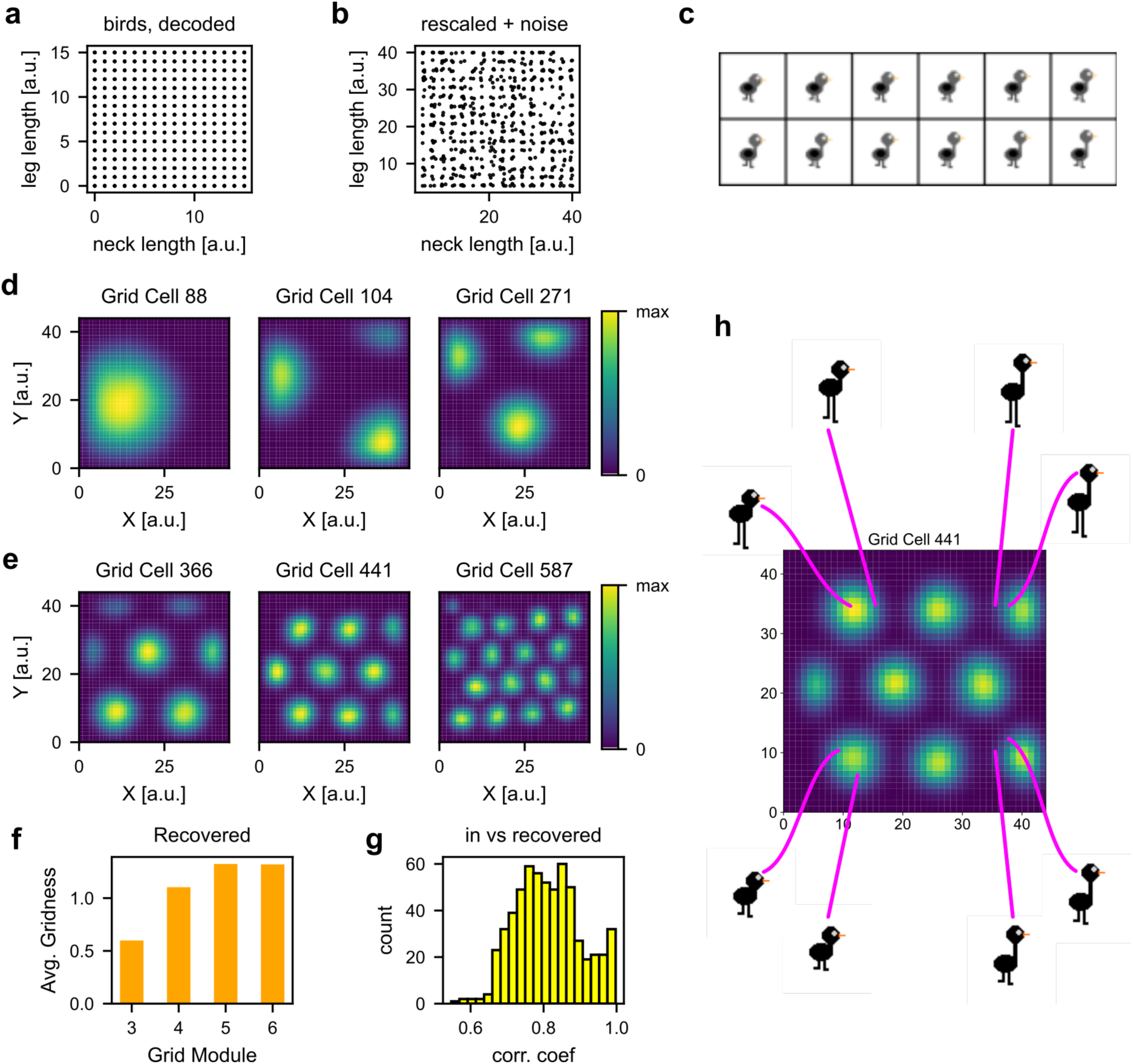
A cognitive map of bird space. **(a)** 1280 bird stimuli are procedurally generated to have a range of leg and neck lengths in discrete classes, and **(b)** noise is added to both properties after rescaling the dataset. **(c)** Bird examples. **(d,e)** Recovered grid cells from all modules. **(f)** Average gridness per module. **(g)** Population histogram of correlation coefficients for correlations between individual recovered grid cells and input grid cells. **(h)** A representative recovered grid cell as a stand-in for the entire stack of *N_GC_* grid cells (cf. population vector illustration in Figure 1a). A pixel corresponds to a grid cell PV. Magenta lines emanate from the locations at which the displayed birds have been anchored. The bird map orders the stimuli according to leg length top to bottom, and neck length left to right.

### Relational Reasoning

What can UCMs be used for^15–17,41^? In the spatial navigation and memory field, it is often assumed that grid cells i) provide a metric for space, ii) perform path integration, and iii) possibly perform vector navigation^26^ (though see ref.^42,43^) in order to calculate a homing vector^44^ or directed trajectories in general. Here we can highlight several types of non-spatial reasoning abilities (relational inference) that can be supported by the present model. Subspace construction, perspective taking and analogical reasoning are explicitly simulated, and additional abilities are outlined in below. Importantly, the key quantities necessary for these reasoning abilities can be calculated without any model additions, though a full neural realization will naturally require additional processing.

Figure 5a shows 4 bird stimuli that were mapped by the PIN to locations near the corners of the grid map (one grid cell rate map shown, representative for the stack of all cells). The model is taken to support analogical reasoning if – given ABC, it can find D such that “C is to D as A is to B” (cf. Figure 5a). Here we can combine the PIN and VNA. Given A and B we can use the VNA to compute the vector AB. Purely for visualization convenience and without loss of generality we pick C horizontally opposite A. Then we engage the PIN with the grid cell PV for point C and the vector AB given by the VNA. This yields a grid cell PV for point D, that stands to C in the same relation as B does to A. If no stimulus is anchored precisely at D, it is assumed that we could pick the nearest. Figure 5b shows 5 simulated examples (for visibility). Pairs of vectors are indicated by similar colors of the lines connecting AB and CD. Note that we could also perform the opposite operation. The VNA could calculate vectors AB and CD to check if an analogy holds (not shown). Figure 5c shows a larger number of simulations. A combination of VNA and PIN can be used to derive analogies on any 2D map that constitutes an arbitrary combination of 2 magnitude codes. Here we could define subsets of data points (e.g., a group of images from the OASIS dataset) and look at average relations/vectors between them (Figure 5c). Such average relations could be interpreted as higher order structure, on top of the relational structure (relative distance and direction) between the individual datapoints/stimuli.

**Figure 5:**
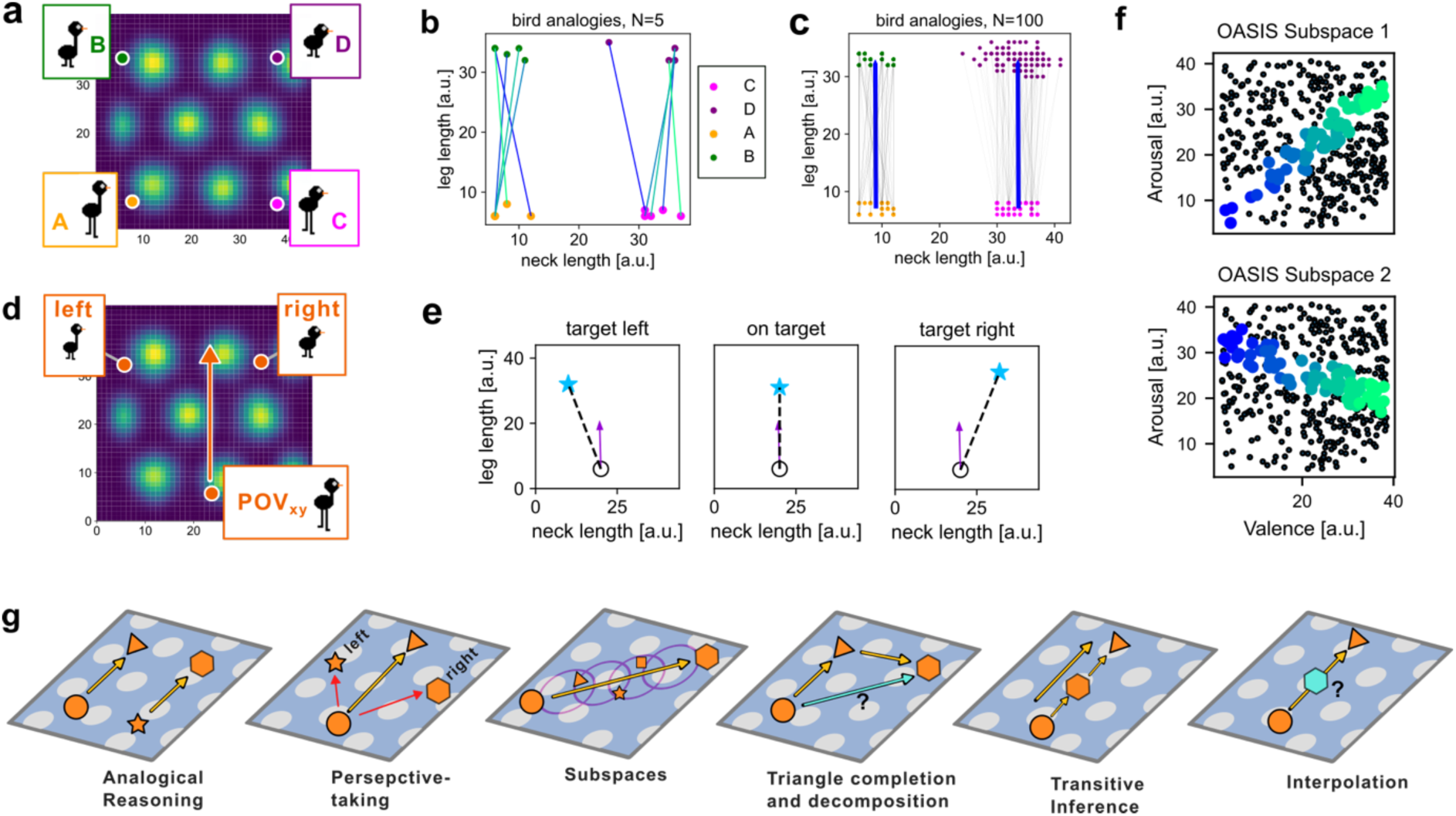
UCMs can facilitate analogical reasoning and other forms of cognition. **(a)** A visual representation of an analogy expressed via the learned cognitive map (here of bird space). A is to B as C is to D, related by the vector ***d***. The VNA can compute ***d*** given A and B. Arbitrarily chosen C with ***d*** put into the PIN yields D. **(b)** 5 analogies (individual simulations connected by same-colored lines) calculated in the model, showing end points on the grid map. **(c)** same as b for 100 analogies. The average relation/vector is superimposed in blue. Relations between task-defined sets could be interpreted as higher order structures. **(d)** If a subject can place him/herself in the abstract map as suggested by ref.^45^ other stimuli could be perceived as lying on the egocentric left or right in abstract space. **(e)** Supplying the model with a “heading” vector and calculating the vector from self to goal allows the model to assign “egocentric-like” bearings to stimuli in abstract space. **(f)** Subspaces can be constructed by finding all points near a connecting vector between two extremes, by scaling the vector in increments and locating nearby points (color changes signal increments). Subspaces can lie on any axis (cf. top and bottom) **(g)** Types of cognition that could be supported by a combination of grid cells, VNA and PIN. Left to right: analogical reasoning (see a-c), egocentric perspective taking (see d), subspace construction (see f), triangle completion (known from spatial navigation research) as a form of relational inference. Transitive inference along an arbitrary direction in 2D as a special case of generalized relational inference (triangle completion). Finally, interpolation: if the cognitive architecture is augmented with a generative network (e.g., the reconstruction part of a variational autoencoder) it could infer a stimulus that is not yet placed on the map.

Next, Viganò and co-workers^45^ showed “egocentric-like” representations in conceptual spaces during goal-directed mental search (also see, ref.^46^). Parietal cortex activity distinguished where a goal was located relative to the subject (e.g., to the left or right) that was mentally navigating on the abstract cognitive map. To compute a signal in the model that can inform this behavior all that is needed is a “heading” vector. The VNA can calculate a vector from self to queried target. Comparing this vector with the heading vector suffices to assign “egocentric-like” bearings (ahead, left, on target) to stimuli in abstract space (Figure 5e). Here this calculation is done simply be computing the signed angle between the two vectors (i.e., not neurally), while the vector to the goal is computed by the VNA. A full neural realization and readout will likely require the involvement of processing outside the hippocampal formation, which, intriguingly, appears to include parietal cortex (precuneus) and retroplenial cortex^45^, much like in models of spatial perspective taking^27,47^.

Both analogy-making and egocentric perspective taking follow a common theme. They result from a set of operations employing grid cells, PIN and VNA in varying combinations that can be executed in sequence to implement acts of cognition. If such a set of operations is independent of the map itself, we can tentatively identify it with a transferable rule set^48^, such as “how to make/check an analogy” or as outlined in Figure 5f,g, subspace construction. In ref.^48^ subjects were tasked to find a 1D subspace defining a category in which two stimulus attributes are of equal relative magnitude (lying on a diagonal). Such a subspace could theoretically be constructed without a cognitive map, though different classification mechanisms are known to compete^49^. The benefit of using UCM-based operations would lie in the ability to apply the set operations on any map (transfer). To build a subspace, more than one set of operations is feasible. We can start with a point and pick a direction (the direction along which the subspace will be spanned), assigning a first vector (e.g., to the far end of the map). The PIN can derive a corresponding grid cell PV at the end of a supplied vector. Even if no stimulus is attached to that PV at the endpoint, we can then search the closest mapped stimulus PVs within some radius and assign it to the subspace. The same vector can then be scaled (shortened or lengthened) in its two components (by the same factor) to locate nearby points at increments along the subspace vector, and so on. Alternatively, the VNA could be used to calculate a vector between two stimuli on the map. Stimuli mapped along this vector can then be retrieved. Figure 5f (top) shows the selected points in the OASIS dataset that have the same valence and arousal on their respective scales. In bird space the diagonal would correspond to all birds where the neck-length is equal to the leg-length. However, here the OASIS dataset is the more useful example, as it illustrates that the two magnitudes (rescaled to their input magnitudes) need not span the same numerical range as they happen to do in bird space. This subspace mapping algorithm generalizes to all angles (Figure 5f, bottom).

Finally, we can sketch out how other types of abstract cognition could be supported by the present model (Figure 5g). It is worth re-iterating that determining the defining quantities that can inform these reasoning abilities does not require additional machinery over and above the present model. That is, the key cognitive operations on the map are supported by the same machinery that was used to build it.

## Discussion

It was shown that the combination of grid cells, VNA and PIN can iteratively build universal cognitive maps under 2 key assumptions: Grid scales are plastic^36,37^, and magnitude codes exist upstream of entorhinal cortex^50^, both of which have support in the literature. Two neurally plausible architectures were suggested for the PIN, while prior work has outlined several implementations for VNAs^24,26^. Any VNA could be substituted here. A common critique of grid cell navigation models is that they rely on perfect hexagonal grids. However, it is not simply a given that distortions impede navigational calculus^43^. Moreover, it was shown that the present model can deal with noise, which simultaneously yields plausible behavioral predictions. Difficult tasks take longer to learn, or may fail completely within allocated time. Importantly, the resultant UCMs can support a variety of reasoning tasks by making use of the very same architectural components that were used to build the maps in the first place. These and other operations could form a composable set that can be applied to any map. If nominally spatial grid cells are repurposed for abstract thought, then spatial navigation deficits go hand in hand with broader cognitive deficits.

The model was tested on two datasets. First, on an experimental dataset (the OASIS dataset^32^), which was also used in the first single cell recordings of non-spatial grid cells^33^. Second, on a procedurally generated version of bird space, similar to the study that provided first, indirect evidence of grid signals for feature maps^9^. Here it is noteworthy that the procedurally generated bird dataset differs from the experiment in that it was sampled in jumps among discrete classes, while the original experiment required subjects to continuously morph the bird stimuli. I.e., a continuum of bird stimuli along navigation trajectories was experienced. Smooth movement of the location estimate on the grid cell manifold by mock motor efference has previously been proposed as a model of mental navigation^27^. The exact same machinery could be used to smoothly travel between grid cell PVs associated with abstract stimuli. As long as mock motor efference translates activity patterns on the grid cells torus, it does not matter that this signal now signifies a “velocity” in abstract space (given a suitable gain setting). The effect on the grid representation remains a translation of the location estimate. This also predicts that participants perceive abstract stimuli on opposite ends of the map as more distant. While smooth motion could be added to the present model, it is not necessary to build a map. Importantly, when confronted with the OASIS dataset, participants did not experience continuous transitions between stimuli. This provides positive proof^33^ that models for the construction of UCMs must be able to cope with discontinuous stimulus sets and cannot solely rely on smooth translation on the grid manifold. The same should hold for visual grid coding when locations are connected by ballistic saccades^38,51^.

Several key open questions remain to be explored. First, how are magnitude codes for given stimulus features extracted from stimuli? Figure 6 proposes a high-level view of the necessary architectural components. The brain must somehow realize which stimulus dimensions are to be mapped^50,52^. A network capable of detecting widely distributed activation of its representational repertoire for a given attribute (e.g., in the early phases of a task) could render that stimulus dimension eligible for mapping, while punctate and sporadic activations along other dimensions signal that a feature is not to be mapped (or cannot be mapped) by the hippocampal-entorhinal system. For example, we can hypothesize that few images in the OASIS dataset showed cars. Hence semantic properties of cars (e.g., size and engine power) are not widely sampled, while – by experimental design – the dataset visits the full gamut of valence and arousal values. Detecting this broad activation could mark those stimulus dimensions for mapping in early task phases. A possible prediction would then be that if more than 2 dimensions vary systematically, the system will choose the 2 dimensions that sample a broader range in early task phases.

**Figure 6:**
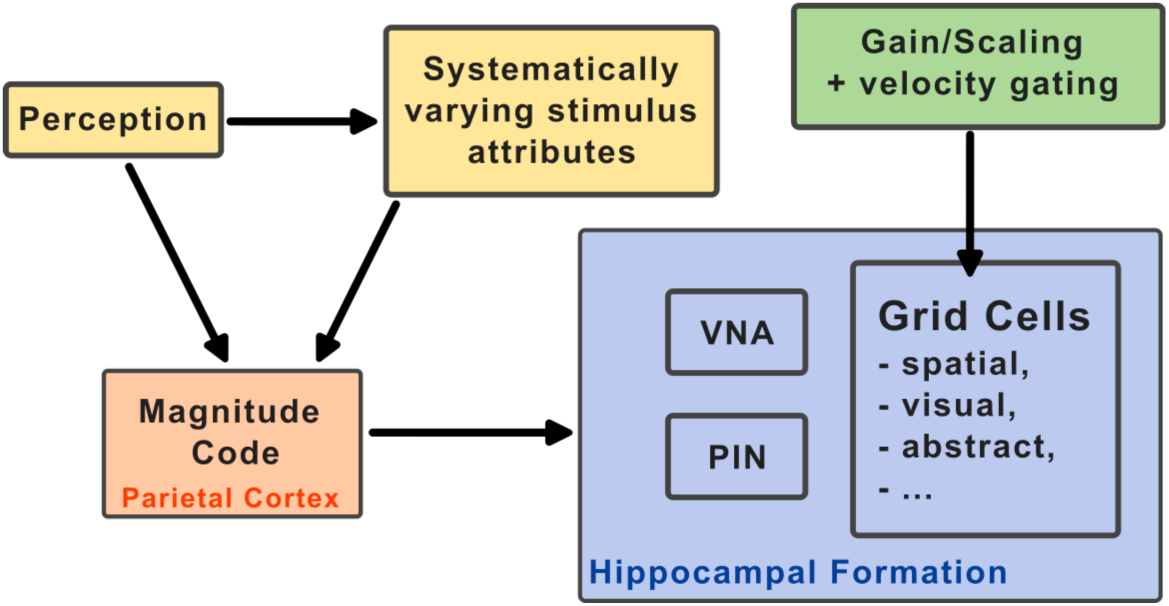
Wider architectural proposal: A VNA, PIN and grid cells must be embedded in a wider cognitive architecture that spans multiple cortical areas, comprised of gating and velocity scaling circuitry (green) to adapt grid cells to a specific task, and a magnitude code (orange) for the individual dimension which are to be mapped onto a UCM. The perceptual system (yellow) must detect which stimulus systems change systematically in a stimulus set, e.g., by detecting widely distributed activation of the representational repertoire for given stimulus features.

Second, how can “structure” transfer between maps^48,55,56^ Figure 5c offered a tentative definition of higher order structure. However, a better candidate for transfer could be a set of operations on the UCM, as outlined in the context of Figure 5. If independent of the map itself, the basic operations form a composable set, and there is an intriguing co-occurrence between rule/task set processing and grid signals in PFC^57,9^.

Third, the notion of landmarks and boundaries in abstract space has previously been identified as a key open question^17^. Here we can speculate that boundary coding^58,59^ could be repurposed to signal the distance to the limits in stimulus magnitudes. Such signals could play a role in the rescaling of grid cells. A domain-general interaction of repurposed boundary codes with grid cells is also suggested by evidence^60^ that could inform updated models of grid coding in the visual domain^38^. The fact that grid-like signals are evident in visual^12,51,60^, social^13,61,62^, auditory^11^, spatial^4^, motor^10^, affective^33^, and abstract^9^ settings, is could mean that the same entorhinal (grid) cells are engaged in all these tasks, rather than disparate, domain-specialized populations. This highlights the need for a domain-general account that is continuous with spatial navigation, allowing for a way to repurpose the existing grid code in novel tasks.

In line with this notion, it was unnecessary here to interpret grid cells as some slowly learned latent variable representation, as seen in machine learning style models^63^. Developmental data^64–66^ suggests there is no need to learn grid cells from scratch^67^ after a given age. That said, a benefit of end-to-end trained models^63,67^ is that connecting actions (between map locations) are explicitly learned. However, in the present model actions are easily derived. Given a VNA we can calculate the connecting vector between any two locations, which directly informs the action on the map (e.g., move up to reach point B) – very much akin to actual spatial navigation. Alternative we could simulate trajectories on the map. Repurposing the navigation system also means that known mechanisms from spatial cognitive maps, such as replay^68^ or remapping^69^ straight-forwardly transfer to UCMs. Post UCM construction, distortions could also be corrected by belief propagation^70^ to adjust an initially imperfect map. The noise simulations shown above suggest that a modest number of replays should suffice. That is, stimuli could be re-anchored offline if pairwise stimulus space distances can be retrieved from memory. Concerning noise, more than one corrective mechanism was proposed above. Additionally, each stimulus could be mapped more than once, to yield average locations and correct initial errors. Similarly, revisiting anchors placed early in the mapping process could correct for drift.

If some form of vector navigation network exists (as suggested by homing in darkness^44^), the PIN could start developing with fully formed vector navigation networks in place, or vice versa. The PIN essentially produces associations between grid cell PVs, gated by distance, and all the entities that are needed for learning (start point, vector, end point) would be available, given a VNA^26^. Alternatively, sampling a sufficient number of random linear trajectories via imagined movement (similar to mental navigation^27^) can generate the necessary quantities. In terms of neural implementations, no extravagant mechanisms were assumed. The PIN would principally rely on synaptic gating. Alternatively, the involvement of head direction cells was suggested. Moreover, the types of relational inference outlined in the context of Figure 5 add important use-cases, in addition to spatial navigation and possibly eye-movement navigation from memory^38^. This makes a vector navigation and inference architecture a highly useful tool in the cortical arsenal of computation and suggests evolutionary pressures to evolve and conserve such an architecture.

In closing, taking the OASIS dataset as an example, it has to be stressed that mapping individual stimuli to locations on a cognitive map of arousal and valence does, of course, not negate the high-dimensional nature of the original stimuli. A UCM is to be viewed as an ad-hoc constructed mapping according to two dimensions that happen to be present in a task. Such maps could be constructed according to task demands and be forgotten as soon as they cease to be relevant, while the mechanisms to build UCMs and operate on them remain in place. The present model shows that this flexibility could in part be accounted for by domain-general spatial navigation architectures applied to non-spatial settings.

## Acknowledgements

AB acknowledges funding from the Max-Planck Society and is grateful for comments on the manuscript by Simone Viganò, Nicholas Menghi, Stephanie Theves, Janis Keck, Viktor Studenyak, and Christian Doeller.

## Extended Methods

Upon acceptance the code to simulate the model will be made available on: https://github.com/Bicanski-NCG

### Grid Cells

Canonical grid cell firing rate maps consist of matrices of dimensions 44 by 44 and are generated by superimposing 60 degree offset cosine waves:

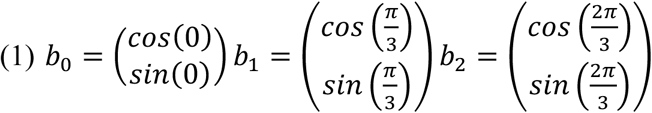

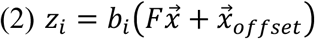

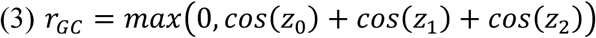

b_i_ are the normal vectors of the waves. 6 modules with varying orientations between them are generated, with the following parameters. 6 modules, with 100 neurons/rate maps each are generated, starting the spatial frequency F = 0.002*2π, increasing approximately by the scaling factor 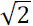 for successive modules^71^. Per module 100 offsets are sampled uniformly, filling the rhomboid of 4 grid vertices on the grid.

### Vector Navigation Architecture (VNA)

The distance-cell model from prior work serves as a VNA^26,72^. Briefly: location in the plane is represented by a grid cell population vector (PV) across all rate maps (dominated by a subset of grid cell with phases that make them fire at the given location). Distances between pairs of grid cells with appropriate PVs can be read out from corresponding distance cells. A total of 4 distance cell arrays is used, 2 for each of two non-co-linear axes. Distance cells project to pairs of readout cells with monotonically increasing/decreasing weights from a given distance cell array. The direction of monotonic increase/decrease of weights is reversed for the other direction along that particular dimension. Relative differences in firing rate between readout neurons represent the vectorial displacement component along the given axis. Note that any VNA could be substituted that operates on the same inputs and yields similar outputs. If one is only interested in the vector, a simpler, albeit less plausible, VNA can be implemented as a lookup-table, taking the current and target grid cell PVs and the stacked GC rate maps to directly return relative distances.

### Positional Inference Network (PIN)

The PIN takes in a grid cell PV and a displacement vector to infer the target PV. Two out of potentially more plausible neural architectures are presented in Figure 1, one based on a distance cell-like model, one based on head direction sweeps. The former is adopted here. Because relative vectors can be identical between two distinct pairs of positions, say, AB, and CD (cf. Figure 1), a given vector displacement **d** connects multiple target grid cell PVs, among which the system has to choose. That is, starting at A and being given **d** the system must output the PV for B, and starting at C and being given the same **d** the system must output the PV for D. The weights for these combinations are pre-calculated and it is assumed that synaptic gating selects the correct weights. In other words, the starting grid cell PV must modulate the selection of the target grid cell PV (cf. Figure 1d). A simplified implementation for the sake of simulation is to have the starting location select a subset of weights, which, when multiplied with a 1-hot distance encoding vector (one out of N elements set to 1, where N corresponds to the range of possible distances on the grid cell rate map along one dimension, i.e., 44.)

### Dataset rescaling and similarity measure

Datasets (Oasis and stretchy birds) are rescaled such that their values lie between 0 and 44 (with some padding, e.g. 4 below), the range of the grid cell rate maps. Example for the Oasis dataset, with valence *v* and arousal *a,* padding *p*, and grid range *res*:

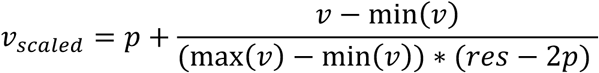

This is to be seen as an approximation of the plastic rescaling of the velocity gain that determines grid scale^20,21,73^, as discussed in the main manuscript. Note, that there is no obligation to use an equal range along both dimensions when rescaling. The vector distance between data points is equated to their similarity. The intuition is that similar stimuli occupy nearby position on the grid map and can thus be treated like spatial navigation distances.

### Anchor and target selection

To assign stimuli (e.g., arousal-valence points or individual birds) to map locations the algorithm begins by placing the central data point of the distribution on the map. Since the grid map is instantiated in the brain as a periodic, toroidal manifold. Hence any starting location is equivalent. Placing the most central data point centrally on the stacked rate maps is simply a convenience in order not to run into the limits of the static rate maps. Then a second data point/stimulus is chosen and a distance along both stimulus dimensions is calculated – yielding the displace vector ***d***. This is always possible since the system has access to the stimuli, even if it does not yet know where to place them on the map. The starting location’s PV and ***d*** are fed into the PIN to yield a goal location grid cell PV. Having placed two stimuli we have 2 anchors for the placement of the next stimulus. From these two anchors the PIN can estimate the target stimulus location and check for overlap (in the simplest version of the model, see next section for multiple comparisons).

### Noise, averaging, and rejection of stimulus placement

As the number of anchors (stimuli placed on the map) grows, the system can make more than 2 distinct estimates of the target location on the grid map (and thus the PV) starting from different anchors. The number of comparisons can be set freely to compensate for the presence of noise. All noise applied to datasets or displacement vector components is Gaussian, with the following standard deviations:

**Table 1:**
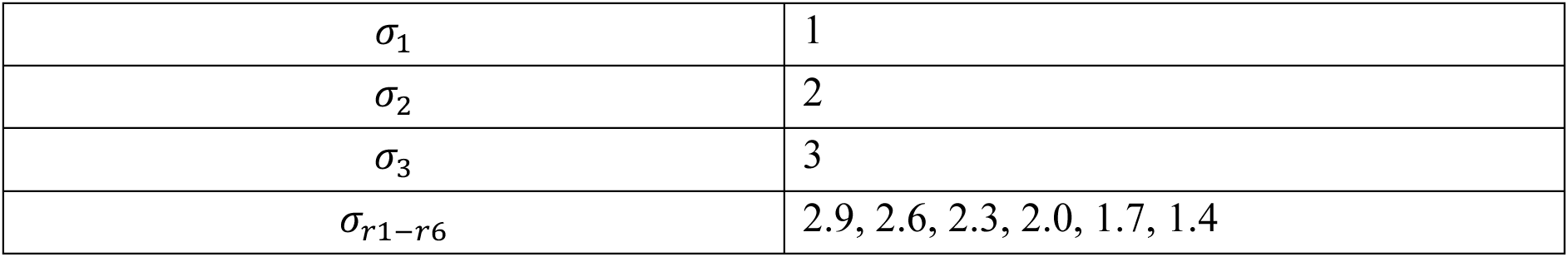
Noise and reconstruction parameters: σ_1-3_ refers to the standard deviations used for low to high vector noise respectively (cf. Figure 3). σ_r1-r6_ refers to the standard deviation of the 2D Gaussian kernel for each module, employed in the grid reconstruction. Picking an intermediate value for all modules yields comparable results.

|  |  |
| --- | --- |
| $\sigma_1$ | 1 |
| $\sigma_2$ | 2 |
| $\sigma_3$ | 3 |
| $\sigma_{r1-r6}$ | 2.9, 2.6, 2.3, 2.0, 1.7, 1.4 |

In the simplest version of the model (no noise) overlap is given if each anchor-target inference via the PIN yields the same coordinates on the grid map starting from 2 anchors, within a small tolerance (here 2 pixels). With more than 2 inferences, an average can be calculated. Since grid cell PVs are read out from stacked rate maps via coordinates on those maps (acting as pointers to grid cell PVs). We can take the median or mean coordinates, or vote on binned coordinates. The median could be preferable if large outliers are expected, and would implicitly implement a kind of first pass error correction. This would be equivalent to putting the inferred target grid cell PV and the starting location grid cell PV into the VNA to check that the resultant vector does not deviate too much from the (dis)similarity in (rescaled) stimulus space. However, in the noise regimes tested here a simply mean across coordinates (pointers) suffices as noise is Gaussian. Coordinates too far from the mean are dropped. If all coordinates lie outside this tolerance the stimulus is considered rejected for map placement. Note that we should not average PVs themselves across coordinates as the output may not correspond to a valid manifold state, assuming attractor dynamics enforce a subset of possible states as valid states.

### Reconstruction of data grids

As the UCM is constructed, inferred PVs are memorized. It is assumed that these grid cell PVs are re-activated by cells that signal the stimulus in the hippocampal-entorhinal system, possibly akin to hippocampal indices. This intuition lets us simulate this re-activation by multiplying the memorized grid cells PV with a place cell-like Gaussian kernel.

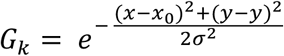

where *x_0_* and *y_0_* are the ground-truth coordinates of the stimulus in the rescaled stimulus space. This is crucial to preserve the potential for mismatch between inferred coordinates (by the PIN) and true coordinates and is akin to an experimenter binning grid cell activity for a given stimulus. Inferred coordinates are only used in the anchor target selection (akin to the model’s internal belief.). We assume that a hippocampal cell was recruited for each stimulus during mapping, which in turn activates the grid cell PV. To reconstruct grid cells from mapped PVs, the PV of each stimulus is replicated across the 44×44 range of the grid maps. The resultant 44×44x*N_GC_* matrix is then multiplied elementwise with the *G_k_* across all rate maps, which squashes activity outside the kernel to zero. This constitutes the contribution of one stimulus to the overall reconstructed rate maps. The procedure is then repeated for all stimuli. Only if the distances on the grid metric respect true stimulus distances will these reconstructions resemble grid cells. Without noise this procedure essentially tests if the PIN correctly navigates across the grid metric. With noise, it tests if additionally, the noise compensation does its job.

### Gridness, autocorrelograms, and correlation coefficients

Gridness is computed in the customary way, established by Hafting et al.^4^. Briefly, for each rate maps (of input grid cells or reconstructed grid cells) the auto-correlogram is calculated (shifting rate maps against each other). Subsequently a ring is cut around the central location in the autocorrelogram and rotated in steps of 3 degrees. For hexagonal grid cells, the symmetry in the autocorrelogram leads too peaks for rotations at 60°, 120°, and 180°, and troughs at 30°, 90° and 150°. The difference between average peaks and troughs is the gridness score. Note that large grid modules with only 1-2 peaks in the rate map show low gridness regardless of perturbation. Hence gridness is complemented by correlation analyses, in which each recovered grid cell is correlated with its counterpart in the set of input grid cells. For the correlation analysis, all recovered grid cells are shifted by the same offset derived from the first stimulus placement on the map vs ground truth coordinates.

### Analogies

Like all simulated relational inferences, to compute analogies, we combine PIN and VNA operations. Given grid cell population vectors for points A and B the VNA to computes the vector AB.

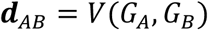

Where *V* signifies the action of VNA, and *G* are grid cell population vectors. Then an arbitrary point C is selected. Purely for convenience we can pick C from a range of possible values on the horizontally opposite side of the map relative to A. Then the PIN is supplied with the grid cell PV for point C and the vector AB given by the VNA. This yields a grid cell PV for point D, that stands to C in the same relation as B does to A.

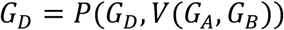

Here *P* signifies the action of the PIN.

### Perspective Taking

To compute egocentric relations to map stimuli we specify a heading vector from an anchor stimulus (our POV position) to a stimulus ahead (*G_Ahead_*, the distance is irrelevant).

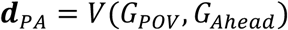

We can then compute the vector to the target that was queried

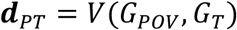

The two vectors ***d****_PA_* and ***d****_PT_* can then be compared to establish egocentric relations, here algorithmically, by computing the signed angle between the two vectors.

### Subspace Construction

To build a subspace, we start with a vector between two extremes, nominally start the end.

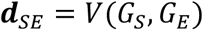

I.e., we pick two stimuli between which we want to span the subspace. This vector is then repeatedly decremented by the same factor to find a grid cell PV.

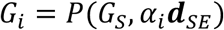

Here *α_i_*<1 is decreased in a coarse-grained manner and a finite number of steps. We then search stimuli with grid cell PVs in a small given radius around *G_i_* on the grid sheet. *G_i_* itself need not be mapped to a stimulus. The found stimuli are tagged as part of the subspace. As an alternative to decrementing the initial vector ***d****_SE_* we can pick a short vector in a given direction and increment *α* > 1.

An complementary procedure for subspace construction is to start with a vector (rather than reference stimuli) and use then PIN to derive a corresponding grid cell PV at the end of a supplied vector and then perform a similar procedure as above. I.e., more than one set of operations of the map could yield a subspace.

## Supplementary Figures

**Figure S1:**
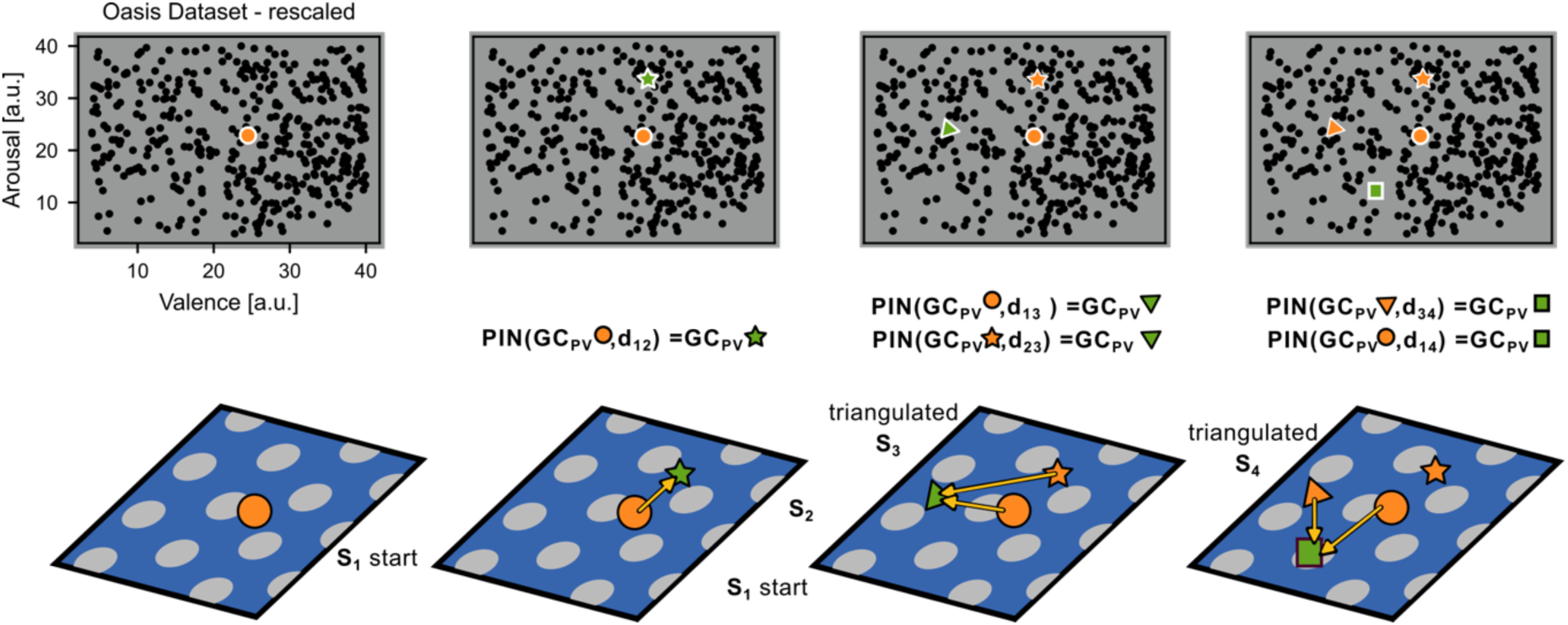
Iterative Mapping - extended illustration. a) Top, dataset with mapped (orange) and to-be-mapped (green) stimuli. Middle row, operation of the PIN network. Bottom row, illustration on a representative grid map. Left-to-right: Mapping the first 4 stimuli. The first stimulus (circle) is mapped at arbitrary location (for simulation convenience near the center). The PIN network infers the grid cell population vector for the green, to-be-mapped, star from the supplied distance (yellow arrow, vector **d**_ij_ in the middle row), placing the star next. As soon as two or more anchors (i.e., already placed stimuli in orange) are present two (and later more) anchors can be queried with respective vectors to the target (yellow arrows) to infer the next target grid cell PV. Multiple anchors introduce the need for agreement, and set up the noise-correction mechanism (cf. Figure 3).

**Figure S2.1:**
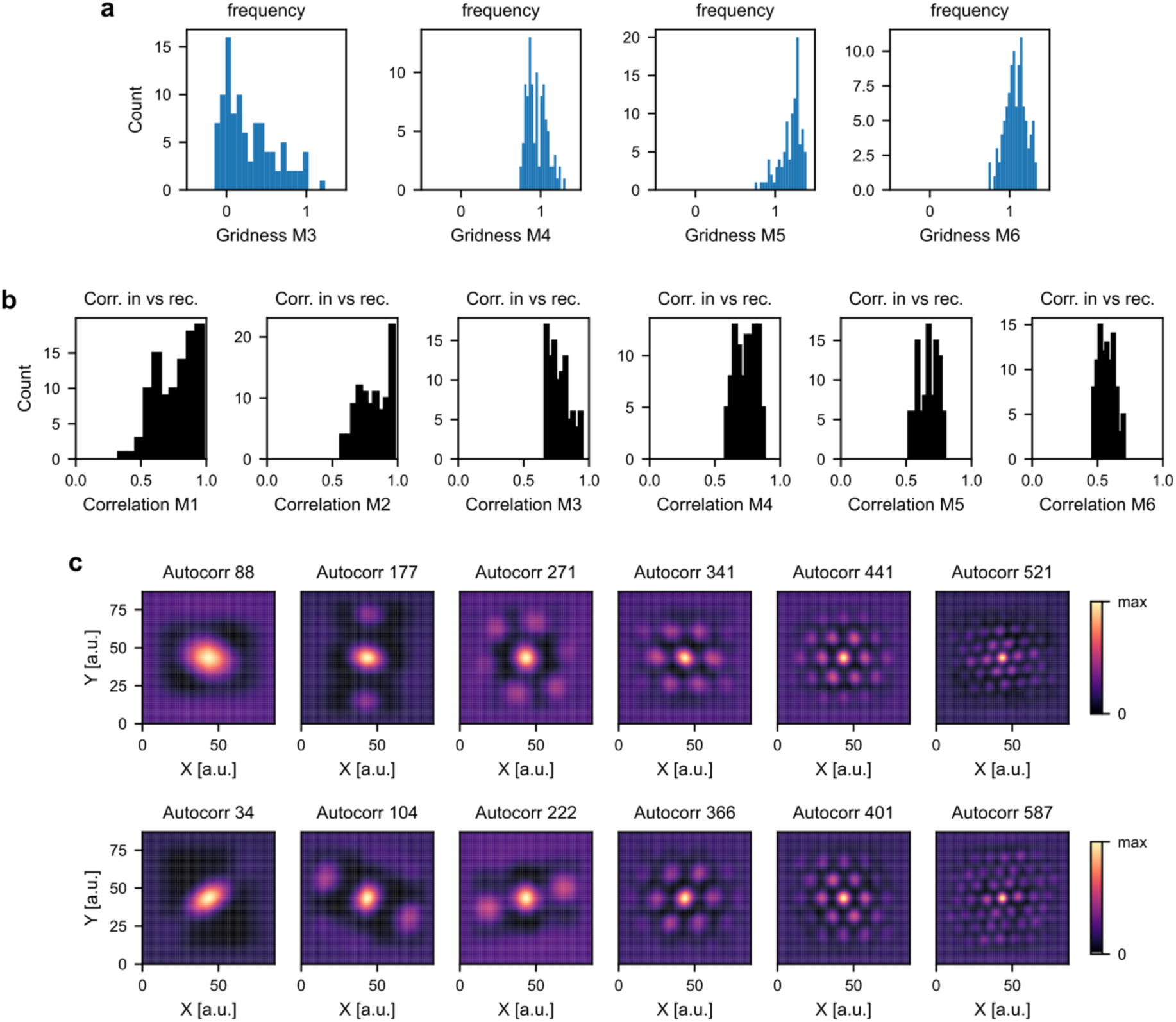
Population histograms and autocorrelograms for the OASIS dataset. a) Histograms of gridness for modules 3 to 6. b) Representative histograms of correlations between input grid cells and recovered grid cells on a cell-by-cell basis. Large scale modules (with only 1-2 peaks per grid map) fail to show high gridness, despite match to input grid cells (cf. Figure 2h) (c) Autocorrelograms of representative grid cells.

**Figure S2.2:**
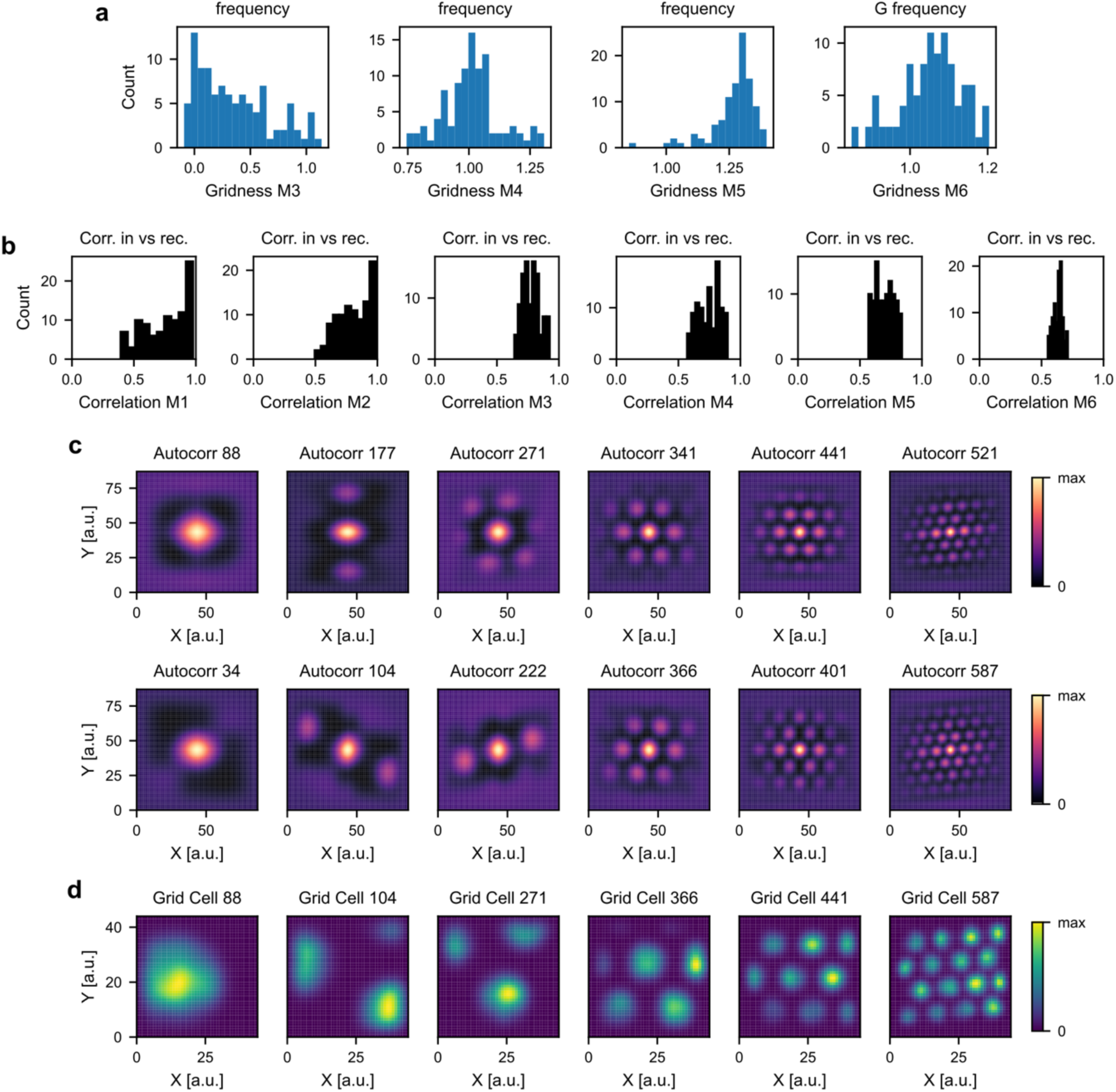
Population histograms and autocorrelograms for synthetic data akin to the OASIS dataset. a) Histograms of gridness for modules 3 to 6. b) Representative histograms of correlations between input grid cells and recovered grid cells on a cell-by-cell basis. Large scale modules (with only 1-2 peaks per grid map) fail to show high gridness, despite match to input grid cells (cf. Figure 2h) (c) Autocorrelograms of representative grid cells. (d) Representative reconstructed grid cells activity maps.

**Figure S3:**
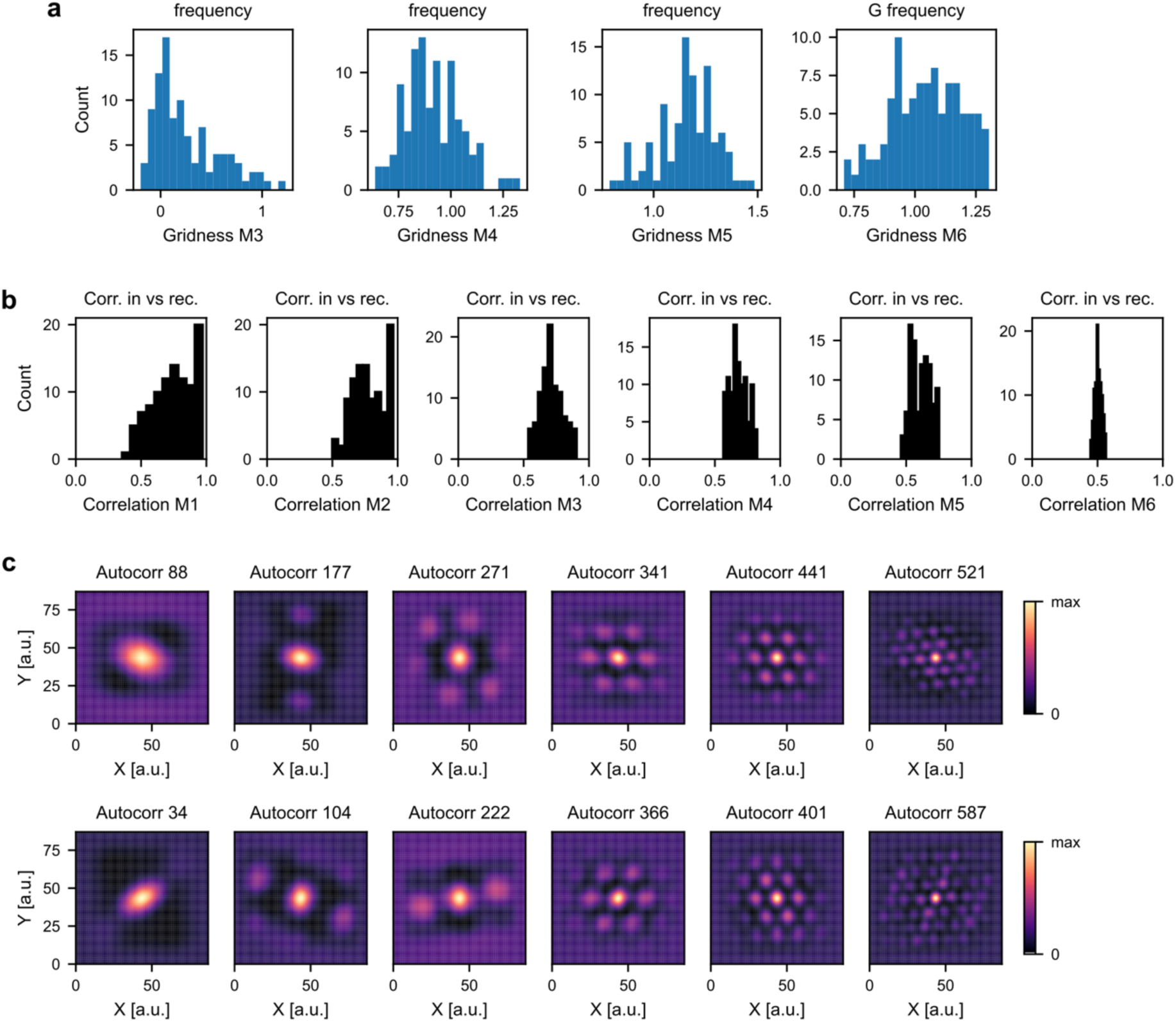
Population histograms and autocorrelograms for the OASIS dataset when noise is added. Averaging recovers a good map, here N=10. a) Histograms of gridness for modules 3 to 6. b) Representative histograms of correlations between input grid cells and recovered grid cells on a cell-by-cell basis. Large scale modules (with only 1-2 peaks per grid map) fail to show high gridness, despite match to input grid cells (cf. Figure 2h) (c) Autocorrelograms of representative grid cells.

